# Accounting for genetic effect heterogeneity in fine-mapping and improving power to detect gene-environment interactions with SharePro

**DOI:** 10.1101/2023.07.27.550862

**Authors:** Wenmin Zhang, Robert Sladek, Yue Li, Hamed S. Najafabadi, Josée Dupuis

## Abstract

**Background:** Characterizing genetic effect heterogeneity across subpopulations with different environmental exposures is useful for identifying exposure-specific pathways, understanding biological mechanisms underlying disease heterogeneity and further pinpointing modifiable risk factors for disease prevention and management. Classical gene-by-environment interaction (GxE) analysis can be used to characterize genetic effect heterogeneity. However, it can have a high multiple testing burden in the context of genome-wide association studies (GWAS) and requires a large sample size to achieve sufficient power.

**Methods:** We adapt a colocalization method, SharePro, to account for effect heterogeneity in finemapping and subsequently improve power for GxE analysis. Through joint fine-mapping of exposure stratified GWAS summary statistics, SharePro can greatly reduce multiple testing burden in GxE analysis.

**Results:** Through extensive simulation studies, we demonstrated that accounting for effect heterogeneity can improve power for fine-mapping. With efficient joint fine-mapping of exposure stratified GWAS summary statistics, SharePro alleviated multiple testing burden in GxE analysis and demonstrated improved power with well-controlled false discovery rate. Through analyses of smoking status stratified GWAS summary statistics, we identified genetic effects on lung function modulated by smoking status mapped to the genes *CHRNA3*, *ADAM19* and *UBR1*. Additionally, using sex stratified GWAS summary statistics, we characterized sex differentiated genetic effects on fat distribution and provided biologically plausible candidates for functional follow-up studies.

**Conclusions:** We have developed an analytical framework to account for effect heterogeneity in finemapping and subsequently improve power for GxE analysis. The SharePro software for GxE analysis is openly available at https://github.com/zhwm/SharePro_gxe.

## 1 Introduction

Genome-wide association studies (GWAS) have led to the identification of numerous genetic loci significantly associated with complex traits and diseases[1–4]. Some genetic loci demonstrate different associations with traits and diseases in subpopulations defined by sex, environmental or lifestyle exposure status [5]. For example, variants in the *NAT2* locus have different associations with bladder cancer in smokers and never smokers [6]. Characterizing such genetic effect heterogeneity can be useful in understanding biological mechanisms underlying disease heterogeneity within a population and further pinpointing modifiable risk factors for prevention and management of complex diseases [7].

Genetic effect heterogeneity can be characterized through gene-by-environment interaction (GxE) analysis. GxE analysis is often performed by including both main effects of a genetic variant and the exposure, as well as a variant-by-exposure interaction term, one variant at a time, in a regression model. Alternatively, in the meta-analysis context, if exposure stratified GWAS summary statistics are available, the heterogeneity test may also be performed to detect GxE [8]. However, due to a high multiple testing burden, a large sample size is usually required to achieve sufficient power to detect GxE in the context of GWAS.

A promising approach for reducing multiple testing burden in the context of GWAS is fine-mapping, a statistical procedure to identify causal variants from GWAS summary statistics and linkage disequilibrium (LD) information [9–11]. With fine-mapped variants, we can effectively reduce the number of tests required in GxE analysis. However, fine-mapping methods usually focus on the inference of causal configurations and do not assess or account for genetic effect heterogeneity, which limited their utility in GxE analysis.

In this work, we adapt SharePro, an accurate and efficient colocalization method that we recently developed [12], to account for effect heterogeneity in fine-mapping and subsequently improve power for GxE analysis. We evaluate the performance of SharePro for fine-mapping in different settings of effect heterogeneity through simulations and compare SharePro with classical methods for GxE analysis in terms of power and false discovery rate (FDR). Furthermore, we examine the utility of SharePro in identifying genetic effects on lung function modulated by smoking status using smoking status stratified GWAS summary statistics. Moreover, we use SharePro to characterize genetic effect heterogeneity in fat distribution with sex stratified GWAS summary statistics.

## 2 Methods

### 2.1 SharePro for GxE analysis

In GxE analysis with SharePro (**Figure 1**), similar to the original SharePro method for colocalization analysis [12] and our sparse projection formulation of the SuSiE model [10, 11, 13], we use a sparse projection to group correlated variants into effect groups. Specifically, in a locus with *G* variants, we assume there are altogether *K* effect groups for traits. We use a sparse indicator **s***_k_* (*k ∈ {*1*, …, K}*) shared across different exposure categories to specify the variant representations for the *k^th^* effect group:

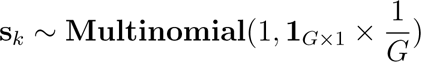

**Figure 1.**
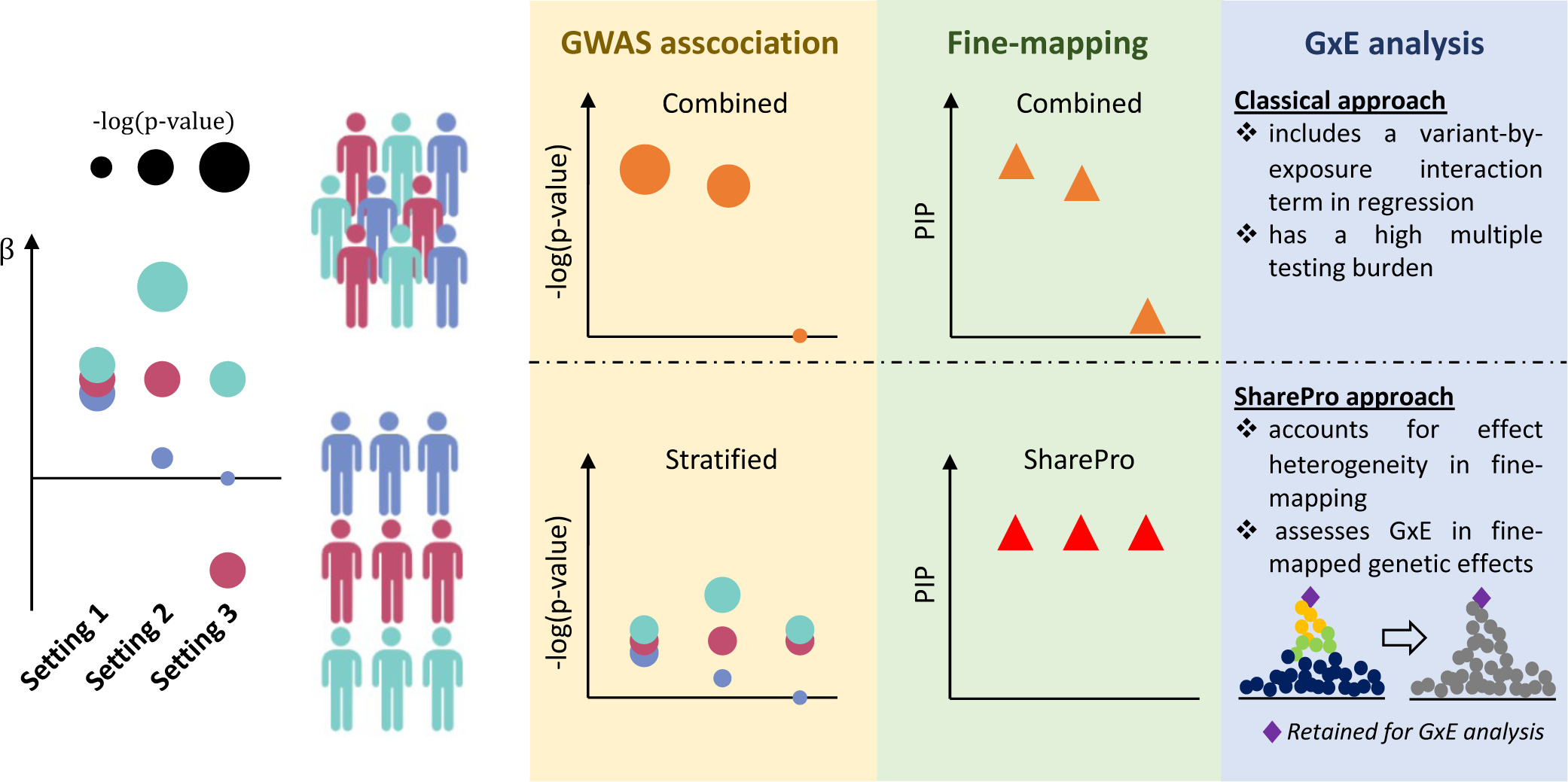
SharePro for GxE analysis. We showcase three settings of effect heterogeneity in a population with three environmental exposure statuses. In setting 1, with similar effect sizes across three exposure categories, this genetic effect can be well-characterized through combined GWAS and fine-mapping of the combined GWAS summary statistics. The stratified approaches might not be well-powered due the relatively small sample sizes in each category. Nevertheless, with SharePro, we can fine-map this genetic effect from exposure stratified GWAS summary statistics and assess its effect heterogeneity. In setting 2, with moderate effect size discrepancy, this genetic effect may be characterized through the combined approaches but the joint approaches will have a higher power. Classical GxE analysis can be underpowered due to a high multiple testing burden. With SharePro, we can first fine-map this genetic effect from exposure stratified GWAS summary statistics and then characterize its effect heterogeneity. With reduced multiple testing burden, we can effectively detect GxE in this setting. In setting 3, the combined approaches can no longer adequately characterize the genetic effect as two exposure categories demonstrates opposite effect sizes. With SharePro, we can accurately fine-map this genetic effect and detect its effect heterogeneity.

Additionally, for each exposure category, we use two sets of exposure specific variables to describe the relationship between the *k^th^* effect group and the trait of interest: *c_ke_* and *c_ku_* for whether the *k^th^* effect group is associated with traits in the exposed and the unexposed category, respectively; *β_ke_* and *β_ku_* for the corresponding effect sizes. Here we illustrate the model with two exposure categories but it is also compatible with multiple exposure categories:

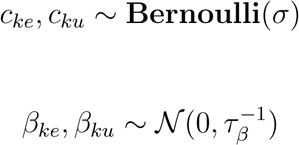

Denoting the genotype matrix in the exposed category as **X***_e_* and the genotype matrix in the unexposed category as **X***_u_*, for traits in the exposed category **y***_e_* and the unexposed category **y***_u_*, we have:

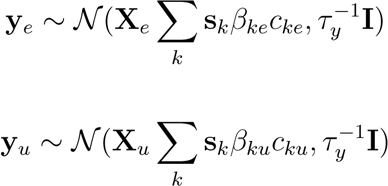

where *τ_β_*and *τ_y_*are hyperparameters for effect size and residual variance; *σ* is the prior probability of effect groups being shared across different exposure categories. We use similar strategies for estimating hyperparameters as in our previous work [11, 12]. With an efficient variational inference algorithm adapted for GWAS summary statistics for posterior inference [12, 14, 15], we can obtain variant-level posterior inclusion probabilities (PIP) for fine-mapping and posterior distributions of effect sizes for GxE analysis. Specifically, the GxE p-value for the *k^th^*effect group can be derived from z-scores:

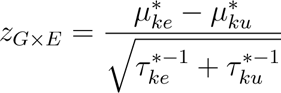

where *µ_ke_^∗^* and *µ_ke_^∗^* are posterior means for effect sizes while *τ_ke_ ^∗−^*^1^ and *τ_ke_ ^∗−^*^1^ are the corresponding posterior variances.

### 2.2 Simulation study design

We conducted simulation studies to examine the benefits of accounting for effect heterogeneity in finemapping and assess the utility of SharePro in GxE analysis. We randomly sampled three 1-Mb region from the genome and extracted genotypes for *N_e_ ∈ {*25000*}* and *N_u_ ∈ {*25000, 10000, 5000*}* non-overlapping UK Biobank European ancestry individuals [1] to simulate traits in the exposed category and the unexposed category, respectively. For each locus, we randomly sampled *K_C_∈ {*1, 2, 3*}* causal variants with effect size *β_e_ ∈ {*0.05*}* in the exposed category and effect size *β_u_ ∈ {−*0.05*, −*0.02, 0, 0.02, 0.05*}* in the unexposed category. The simulation process was repeated 50 times for each setting.

### 2.3 Association test and fine-mapping in simulation studies

We first used PLINK [16] to perform a GWAS separately in the exposed category and the unexposed category and obtained exposure stratified GWAS summary statistics for each simulated trait. We also performed regression analyses using a popular method, GEM [17], with individual-level genotype and exposure status to obtain results for the 1-df association test, the 2-df association test and the GxE test. Specifically, the 1-df association test assesses the marginal variant-trait associations, combining data from all exposure categories. The GxE test assesses whether variant-trait associations differ across exposure categories. The 2-df association test combines the 1-df association test and the GxE test and jointly assesses variant-trait associations and association differences across exposure categories [18].

We also applied SharePro with the exposure stratified GWAS summary statistics and in-sample LD calculated by PLINK to obtain joint fine-mapping results. Additionally, we performed combined finemapping with the 1-df association test GWAS summary statistics and in-sample LD using SharePro. We compared the area under precision-recall curve (AUPRC) in identifying simulated causal variants calculated from SharePro fine-mapping PIP, combined fine-mapping PIP, the 1-df association test p-values and the 2-df association test p-values.

### 2.4 GxE analysis in simulation studies

We obtained effect group-level GxE p-values from SharePro. For the ease of comparison with existing variant-level GxE analysis methods, we derived variant-level GxE p-values for SharePro by assigning effect group-level GxE p-values to variants included in effect groups.

We compared SharePro effect group-level GxE p-values (SharePro effect), and SharePro GxE pvalues mapped to variants in effect groups (SharePro variant) with GEM GxE p-values calculated from individual-level data, Bonferroni corrected GEM GxE p-values (GEM-Bonferroni), Benjamini-Hochberg (BH) corrected GEM GxE p-values (GEM-BH) and GxE p-value derived from the heterogeneity test using exposure stratified summary statistics from PLINK in terms of power and false discovery rate (FDR) at a p-value cutoff of 0.01.

### 2.5 Gene-by-smoking interactions in lung function

Smoking is one of the most common risk factors for chronic obstructive pulmonary disease and has a significant impact on lung function [19]. We investigated genetic effects on lung function that may be modulated by smoking status. Specifically, we focused on analyzing the ratio of the forced expiratory volume in the first one second to the forced vital capacity of the lung measured by spirometry (FFR), a widely used biomarker for lung function.

We performed self-reported smoking status stratified GWAS for FFR in individuals of European ancestry in the UK Biobank [1], including 27,409 current smokers, 100,285 past smokers and 155,951 never smokers. FFR measurements were inverse rank normal transformed after adjusting for age, agesquared, sex, genotyping array, recruitment center, and the first 20 genetic principal components.

We applied SharePro to these smoking status stratified GWAS summary statistic with the UK Biobank European ancestry LD matrix calculated by Weissbrod et al [20]. Effect groups with a gene-by-smoking interactions (GxSmoking) p-value smaller than 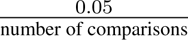 were considered effect group-level Bonferroni significant. We annotated each effect group to the nearest protein coding gene based on UCSC genome annotations [21]. Additionally, we performed the heterogeneity test with smoking status stratified GWAS summary statistics and obtained variant-level GxSmoking p-values.

### 2.6 Gene-by-sex interactions in fat distribution

Fat distribution has important implications for health and is associated with metabolic and cadiovascular diseases [22–24]. We investigated sex differentiated genetic effects on body fat distribution, measured by waist-to-hip ratio adjusted for body mass index (WHRadjBMI). We performed sex stratified GWAS for WHRadjBMI in individuals of European ancestry in the UK Biobank [1], including 189,830 females and 162,508 males. The waist-to-hip ratio measurements were inverse rank normal transformed after adjusting for body mass index, age, age-squared, sex, genotyping array, recruitment center, and the first 20 genetic principal components. We applied SharePro to the sex stratified GWAS summary statistics with the UK Biobank European ancestry LD matrix calculated by Weissbrod et al [20]. Effect groups with a gene-by-sex interactions (GxSex) p-value smaller than 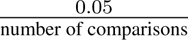 were considered effect group-level Bonferroni significant. We annotated each effect group to the nearest protein coding gene based on UCSC genome annotations [21].

Next, we investigated whether GxSex in WHRadjBMI may be explained by GxSex in sex hormones and lipid metabolism. We obtained sex stratified GWAS summary statistics for bioavailable testosterone and sex hormone binding globulin (SHBG) for UK Biobank European ancestry from Ruth et al [25]. Additionally, we performed sex stratified GWAS using European ancestry individuals in the UK Biobank for triglycerides (TG), low density lipoprotein cholesterol (LDL) and high density lipoprotein cholesterol (HDL). All measurements were inverse rank normal transformed after adjusting for age, age-squared, sex, genotyping array, recruitment center, and the first 20 genetic principal components.

Additionally, we examined whether variants in effect groups exhibited different minor allele frequencies (MAF) in females and males, which could indicate sex differentiated participation bias [26]. We obtained p-values for sex differentiated MAF from z-scores:

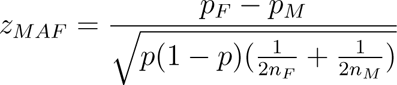

where *p_F_*and *p_M_* are MAF in females and males while *n_F_*and *n_M_* correspond to sample sizes in females and males and *p* is the overall MAF.

## 3 Results

### 3.1 Accounting for effect heterogeneity improved power for fine-mapping

To examine the benefits of accounting for effect heterogeneity in fine-mapping, we conducted simulation studies varying sample sizes (*N_e_* and *N_u_*), effect sizes (*β_e_* and *β_u_*) and number of causal variants (*K_C_*). As expected, in the presence of simulated effect heterogeneity (*β_e_* ≠ *β_u_*), the 2-df association test, which accounts for effect heterogeneity, exhibited a higher AUPRC compared to the 1-df association test. SharePro fine-mapping accounted for both effect heterogeneity and LD, thereby achieving the highest AUPRC. For example, with three causal variants having an opposite effect sizes in the exposed and the unexposed category of equal sample size (*N_e_* = *N_u_* = 25, 000; *β_e_* = 0.05; *β_u_* = *−*0.05 and *K_C_* = 3), the 1-df association test and the combined fine-mapping both had an AUPRC *<* 0.01 (**Figure 2** and **Supplementary Table** S1), because variant-trait associations in the overall population were not detectable. In contrast, the 2-df association test achieved a median AUPRC of 0.34, while SharePro finemapping achieved a median AUPRC of 0.96 (**Figure 2** and **Supplementary Table** S1).

**Figure 2.**
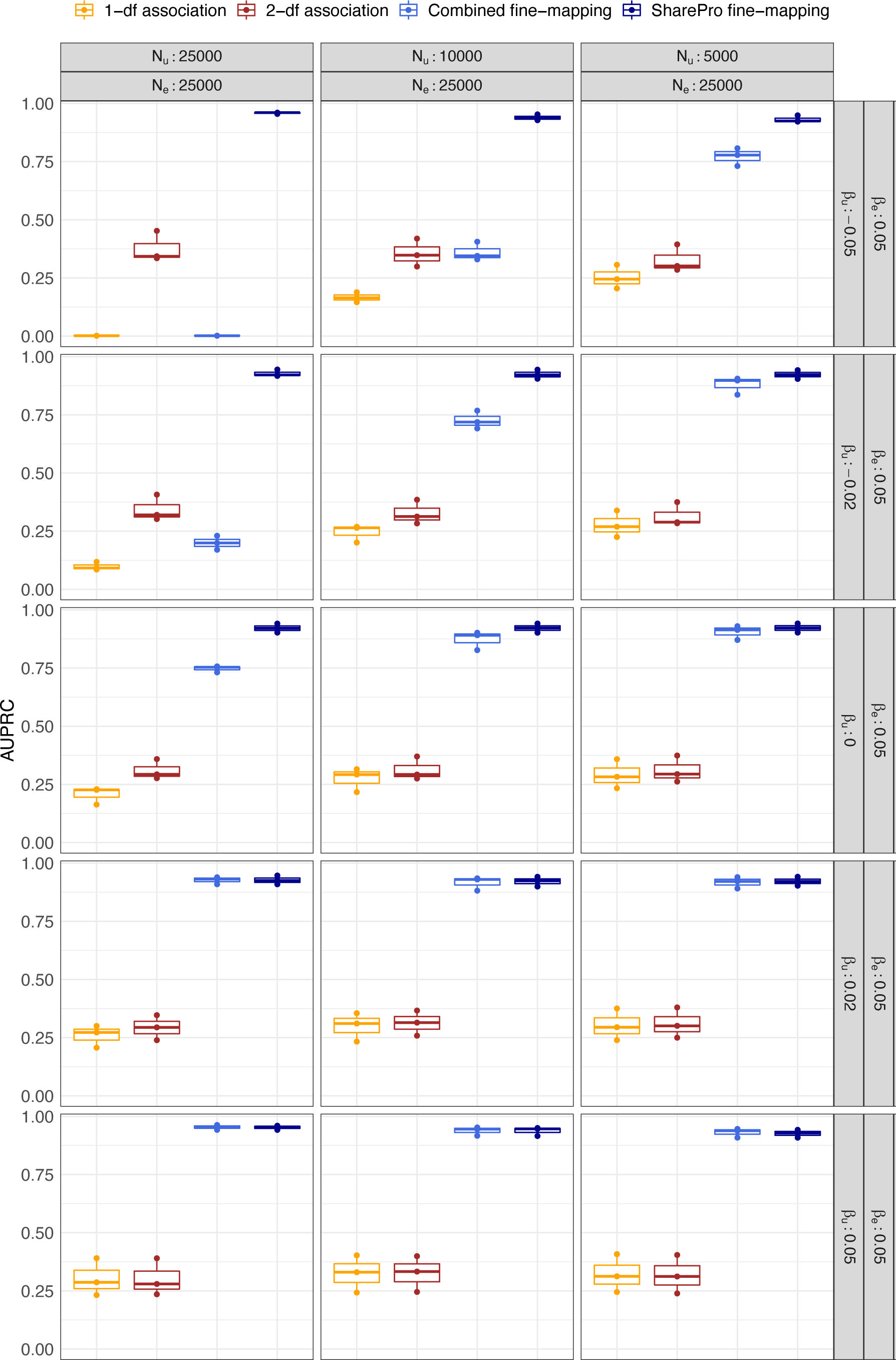
Accounting for effect heterogeneity improved power for fine-mapping. Area under precision and recall curves (AUPRC) in simulation settings with 3 causal variants in a locus (*K_C_* = 3) are displayed. Rows represent different effect size discrepancies with *β_e_*and *β_u_*correspond to effect size in the exposed and the unexposed category. Columns represent different sample size settings with *N_e_* and *N_u_* correspond to sample sizes in the exposed and the unexposed category. Median AUPRC are indicated by horizontal bars and inter-quartile ranges are represented by boxes.

As the magnitude of the effect size discrepancy (*β_e_ − β_u_*) decreased, the performance differences between methods using the joint approaches and those employing the combined approaches attenuated.

For example, with causal variants having a effect size discrepancy of 0.07 (*β_e_* = 0.05; *β_u_* = *−*0.02; *N_e_* = *N_u_* = 25, 000 and *K_C_* = 3), SharePro fine-mapping had a median AUPRC of 0.92 while combined fine-mapping had a median AUPRC of 0.20 (**Figure 2** and **Supplementary Table** S1). Meanwhile, 2-df association test achieved a median AUPRC of 0.32 and 1-df association test only had a median AUPRC of 0.09 (**Figure 2** and **Supplementary Table** S1). When the effect size difference decreased to 0.05 (*β_e_* = 0.05; *β_u_* = 0.00), SharePro fine-mapping maintained a median AUPRC of 0.92 and combined fine-mapping achieved a median AUPRC of 0.75 (**Figure 2** and **Supplementary Table** S1).

Similarly, as we decreased effect heterogeneity by reducing the sample size of the unexposed category, the performance difference also became smaller. For example, with three causal variants having an opposite effect in the exposed category and the unexposed category of unequal sample sizes (*N_e_* = 25, 000; *N_u_* = 10, 000; *β_e_* = 0.05; *β_u_* = *−*0.05 and *K_C_* = 3), the median AUPRC of SharePro fine-mapping was 0.94 while the median AUPRC of combined fine-mapping was 0.34 (**Figure 2** and **Supplementary Table** S1). With a smaller sample size for the unexposed category (*N_u_* = 5, 000), the median AUPRC of SharePro fine-mapping was 0.92 while the median AUPRC of combined fine-mapping was 0.78 (**Figure 2** and **Supplementary Table** S1).

The results for simulations with one or two causal variants exhibited similar patterns and are detailed in **Supplementary Table** S1.

### 3.2 SharePro demonstrated improved power with controlled FDR in GxE analysis

We next examined the utility of SharePro in GxE analysis and compared it to GEM and the heterogeneity test implemented in PLINK using exposure stratified summary statistics. SharePro consistently achieved the highest power with the lowest FDR across different simulation settings. For example, when the effect size discrepancy (*β_e_ − β_u_*) was large (*β_e_*= 0.05, *β_u_* = *−*0.05 and *K_C_* = 3), at a p-value cutoff of 0.01, all methods achieved a similar power close to 1 (**Figure 3A** and **Supplementary Table** S2). However, with the most stringent multiple testing correction, GEM-Bonferroni still had a mean FDR of 0.89 (**Figure 3A** and **Supplementary Table** S2). In contrast, the SharePro effect-group level GxE p-value had the lowest mean FDR of 0.01 while the SharePro GxE p-value mapped to variants in effect groups (SharePro variant) had a mean FDR of 0.33 (**Figure 3A** and **Supplementary Table** S2).

**Figure 3.**
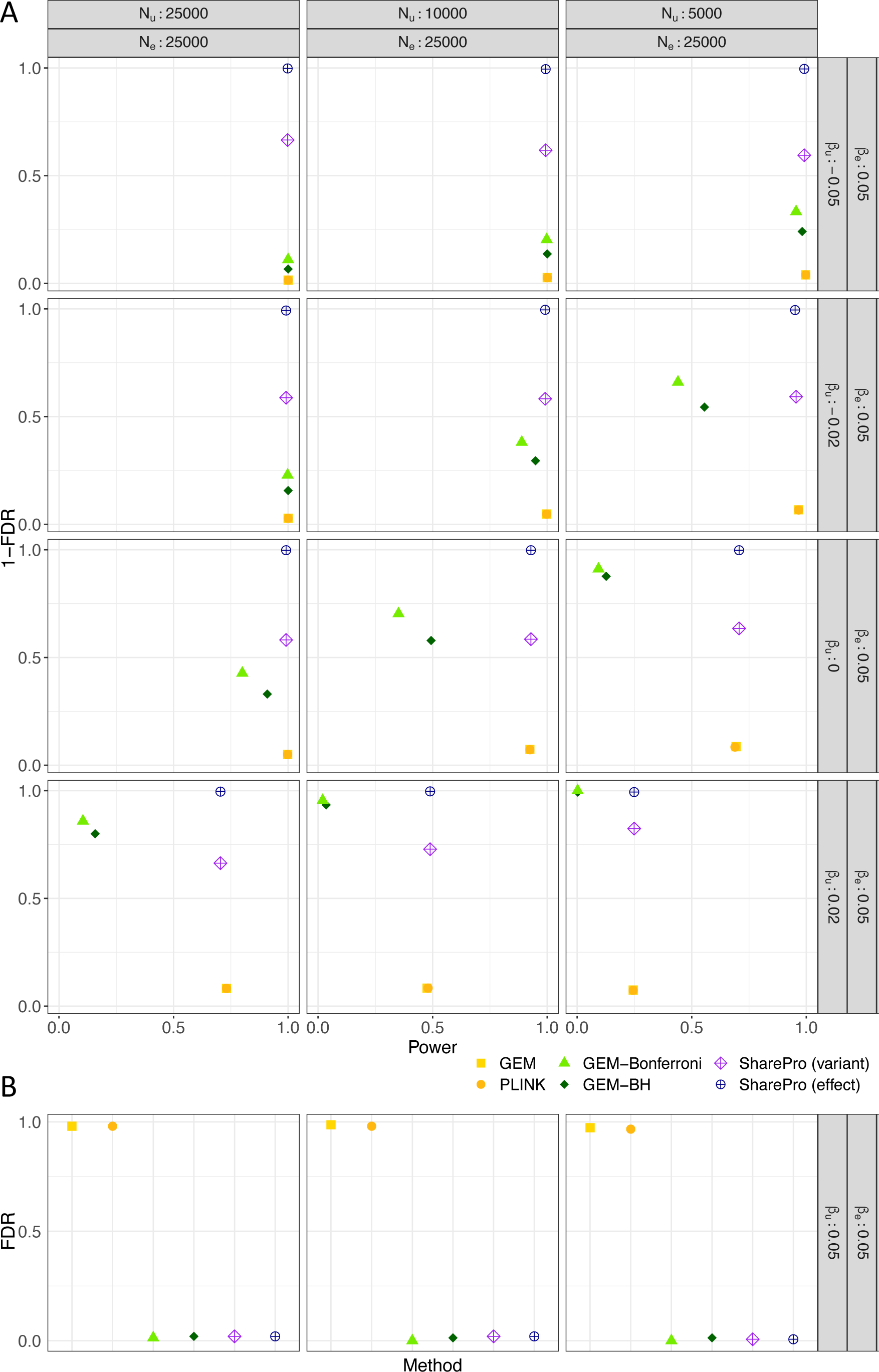
SharePro demonstrated improved power with controlled false discovery rate (FDR) in GxE analysis. (A) Power and 1-FDR in GxE analysis in heterogeneous settings with a p-value cutoff of 0.01. The x-axis stands for power and the y-axis stands for 1-FDR and each dot represents a GxE analysis method. (B) FDR in GxE analysis in homogeneous settings at a p-value cutoff of 0.01. The x-axis stands for methods and the y-axis stands for FDR. As expected, the GxE p-values calculated by GEM using individual-level data and GxE p-values calculated by the heterogeneity test using stratified summary statistics from PLINK were highly similar.

With smaller effect size discrepancy (*β_e_ − β_u_*), SharePro maintained a high power with a low FDR, whereas other methods had decreased power. For instance, in the simulation setting with three causal variants having a effect size discrepancy of 0.03 between the exposed and the unexposed category of equal sample size (*β_e_* = 0.05; *β_u_* = 0.02; *N_e_* = *N_u_* = 25, 000 and *K_C_* = 3) (**Figure 3A** and **Supplementary Table** S2), SharePro achieved an mean power of 0.70 with an mean FDR of 0.35 at the variant-level and a mean FDR lower than 0.01 at the effect group-level. After multiple testing correction, GEMBH and GEM-Bonferroni achieved an mean power of 0.16 and 0.10 with an mean FDR of 0.20 and 0.14, respectively. Notably, without effect heterogeneity (*β_e_* = *β_u_* = 0.05) (**Figure 3B**), both SharePro and classical GxE methods corrected for multiple testing demonstrated a well-controlled FDR.

SharePro differs from classical GxE methods in that it uses joint fine-mapping to reduce the multiple testing burden. We illustrate the importance of this difference with a simulation example (*β_e_* = 0.05; *β_u_* = 0.02; *N_e_* = *N_u_* = 25, 000, *K_C_* = 1) shown in **Figure 4**. In this example, the association test in the exposed and the unexposed categories exhibited clear differences (**Figure 4A**) and the GxE test in GEM accurately detected the simulated effect heterogeneity (**Figure 4B**). However, the power for this test diminished after adjusting for multiple testing (**Figure 4B**). In contrast, SharePro identified a candidate effect group through joint fine-mapping (**Figure 4C**), thus avoiding testing on every variant, resulting in a successful GxE detection (**Figure 4D**).

**Figure 4.**
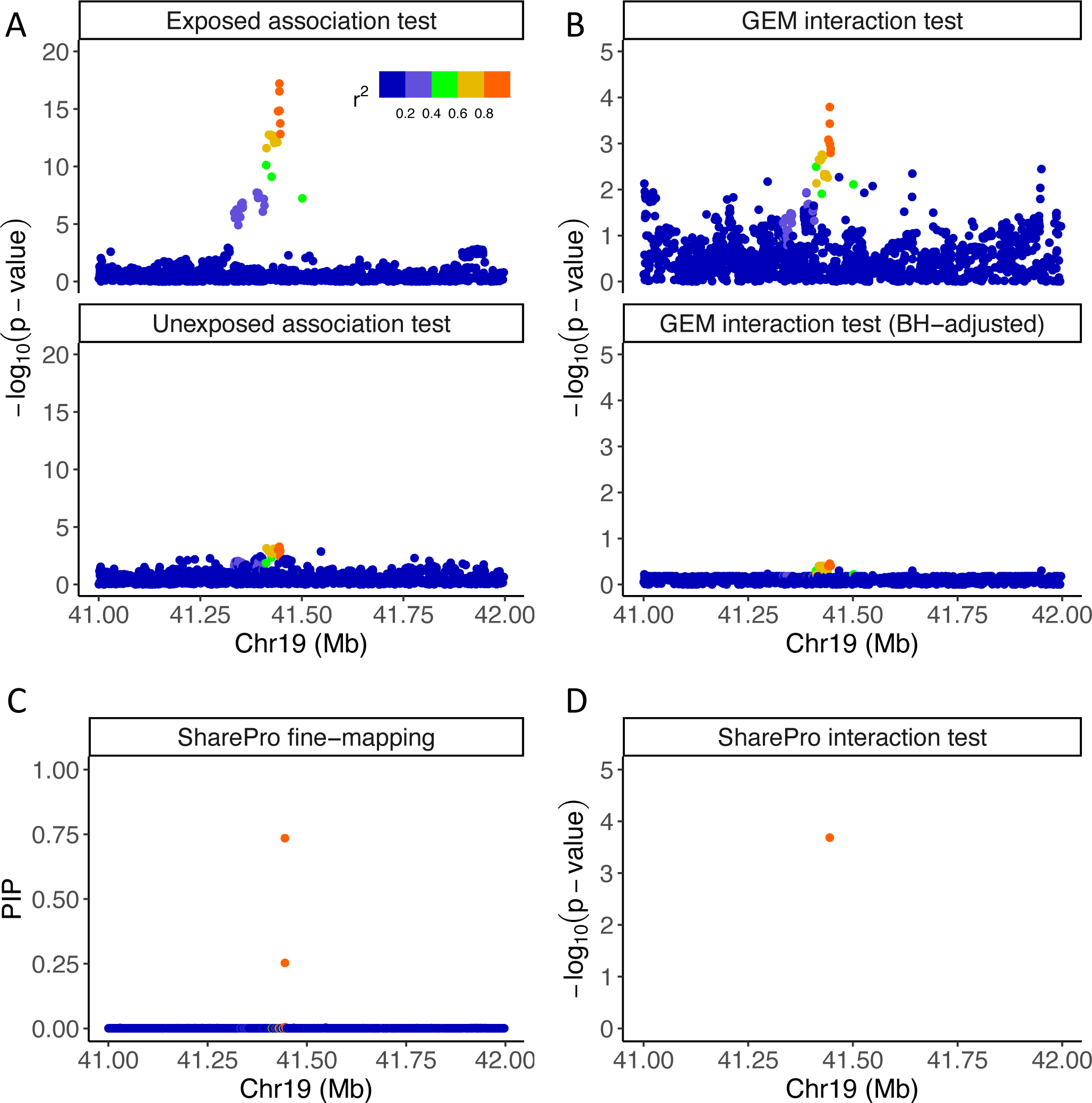
SharePro reduced multiple testing burden in GxE analysis. (A) Exposure stratified GWAS summary statistics in one simulation setting (*β_e_* = 0.05; *β_u_* = 0.02; *N_e_* = *N_u_* = 25, 000, *K_C_* = 1). Each dot represents a variant and the color indicates its correlation with the simulated causal variant. (B) GEM GxE p-values and BH-adjusted GEM GxE p-values. After multiple testing correction, classical GxE analysis had insufficient power to detect effect heterogeneity. (C) Posterior inclusion probabilities in SharePro fine-mapping. Two variants in high LD were identified in an effect group. (D) The GxE p-value for the effect group identified in SharePro fine-mapping.

Moreover, SharePro also demonstrated high computational efficiency. Across different simulation settings, on average, PLINK took 1.0 second to compute stratified GWAS summary statistics (**Supplementary Table** S3), which were used as input for SharePro. SharePro then took 4.5 seconds to perform GxE analysis while GEM on average took 5.8 seconds to calculate test statistics (**Supplementary Table** S3).

### 3.3 SharePro identified genetic effects on lung function modulated by smoking status

Based on smoking status stratified GWAS for FFR, we investigated genetic effects on lung function modulated by smoking status. Across current smokers, past smokers and never smokers, we identified altogether 298 effect groups associated with lung function (**Supplementary Table** S4). Significant GxSmoking were observed in the effect group mapped to the *CHRNA3* gene, comparing current smokers with past smokers (**Figure 5A**), current smokers with never smokers (**Figure 5B**), and past smokers with never smokers (**Figure 5C**). Variants in this effect group also demonstrated genome-wide (5.0 *×* 10*^−^*^8^) significance in the heterogeneity test comparing current smokers with never smokers (**Supplementary Figure** S1B), and past smokers with never smokers (**Supplementary Figure** S1C) while the difference between current smokers and past smokers was close to be significant (**Supplementary Figure** S1A).

**Figure 5.**
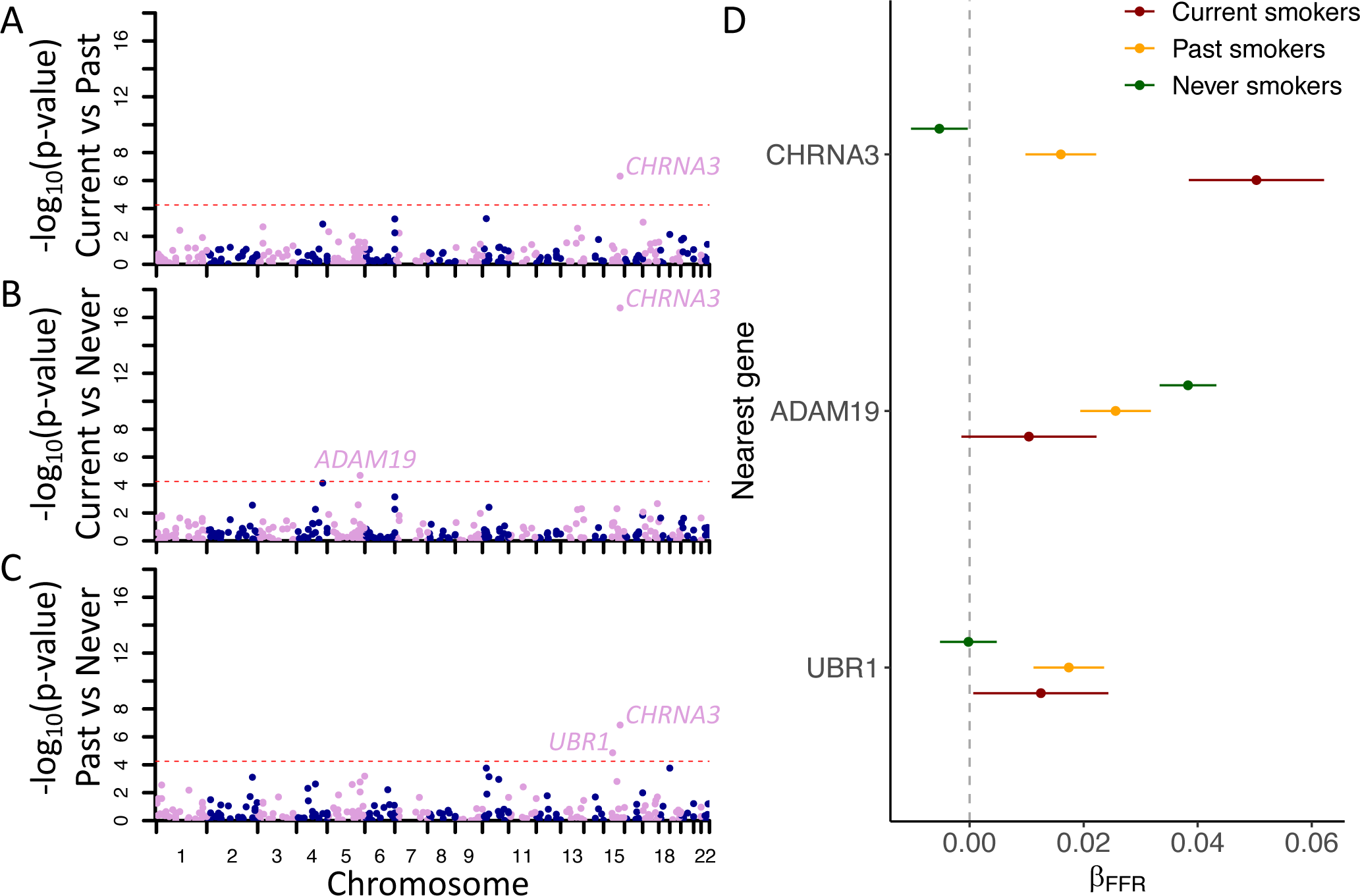
SharePro identified genetic effects on lung function modulated by smoking status. Effect group-level gene-by-smoking interactions (GxSmoking) p-values comparing (A) current smokers with past smokers (B) current smokers with never smokers (C) past smokers with never smokers. The red dotted line corresponds to the effect group-level Bonferroni significant threshold. (D) Effect sizes of effect group-level Bonferroni significant GxSmoking. The x-axis stands for posterior mean effect sizes and the colors represents different smoking statuses.

Additionally, the effect group mapped to the *ADAM19* gene demonstrated effect group-level Bonferroni significant GxSmoking between current smokers and never smokers (**Figure 5B**) while the effect group mapped to the *UBR1* gene demonstrated effect group-level Bonferroni significant GxSmoking between past smokers and never smokers (**Figure 5C**).

Interestingly, the estimated effect sizes of these effect groups across different smoking statuses suggest potential mechanisms of GxSmoking. The effect group mapped to the *CHRNA3* gene demonstrated a large effect size in current smokers, a moderate effect size in past smokers and a minimal effect size in never smokers (**Figure 5D**). The *CHRNA3* gene is known to play a significant role in nicotine dependence and smoking behaviors [27–30]. Our results indicated that the genetic effect of *CHRNA3* on lung function may be mediated by smoking, aligning with its biological function.

Moreover, the effect group mapped to the *ADAM19* gene demonstrated an opposite trend with a large effect size in never smokers and a small effect size in current smokers (**Figure 5D**). The ADAM (a disintegrin and metalloprotease) family has been implicated in regulation of airway inflammation [31, 32]. Further investigations are necessary to confirm whether smoking-induced inflammation may disrupt such regulatory roles. The effect group mapped to the *UBR1* gene demonstrated comparable effect sizes in current smokers and past smokers and no effect in never smokers (**Figure 5D**). *UBR1* encodes a ubiquitin ligase that plays an important role in modulating MGMT (a DNA repair enzyme) homeostasis [33], encouraging future investigations into its physiological role in repairing smoking-induced tissue damages.

### 3.4 SharePro improved characterization of sex differentiated genetic effects on fat distribution

Based on sex stratified GWAS for WHRadjBMI, we investigated sex differentiated genetic effects on fat distribution and identified altogether 293 effect groups associated with WHRadjBMI (**Supplementary Table** S5). Of these, 89 effect groups demonstrated effect group-level Bonferroni significant GxSex and among them 81 effect groups demonstrated a larger effect size in females (**Supplementary Table** S5).

Effect groups demonstrated significant GxSex were mapped to genes that play important roles in adipose cell biology (**Supplementary Table** S5). For example, *COBLL1* has been shown to play a crucial role in actin cytoskeleton remodeling during the differentiation of metabolically active and insulin-sensitive adipocytes [34] and has been observed to exhibit sex dimorphic expression [34]. Recent research indicates RSPO3 can impact body fat distribution by regulating the expansion of gluteofemoral adipose tissue [35] while estrogen is a key regulator of adipogenesis for gluteofemoral adipocytes [36–38]. Additionally, KLF14 is a transcription factor that regulates gene expression in adipose tissue [39]. Recent studies have demonstrated that *KLF14* can increase adiposity and redistribute lipid storage in female mice, but not in male mice [40]. Moreover, SharePro identified an effect group with one variant rs1534696, mapped to the *SNX10* gene, demonstrating significant GxSex in WHRadjBMI (p-value = 6.2 *×* 10*^−^*^11^; **Supplementary Table** S5). This variant has been shown to regulate diet-induced adipose expansion in female mice, but not in male mice [41].

We then assessed whether these sex differentiated genetic effects could be partially attributed to known factors of sex differences, including sex hormones and lipid metabolism. We observed that the top variants from effect groups demonstrating GxSex in WHRadjBMI also tend to have sex differentiated effects on SHBG level (Spearman correlation = 0.43; p-value = 1.6 *×* 10*^−^*^14^; **Figure 6A**). Interestingly, effect groups exhibiting effect group-level GxSex in WHRadjBMI that did not reach genome-wide significance also showed a large effect size discrepancy on SHBG between females and males, such as those mapped to *MAFB*, *FAM13A*, and *TRIB1* (**Figure 6A**).

**Figure 6.**
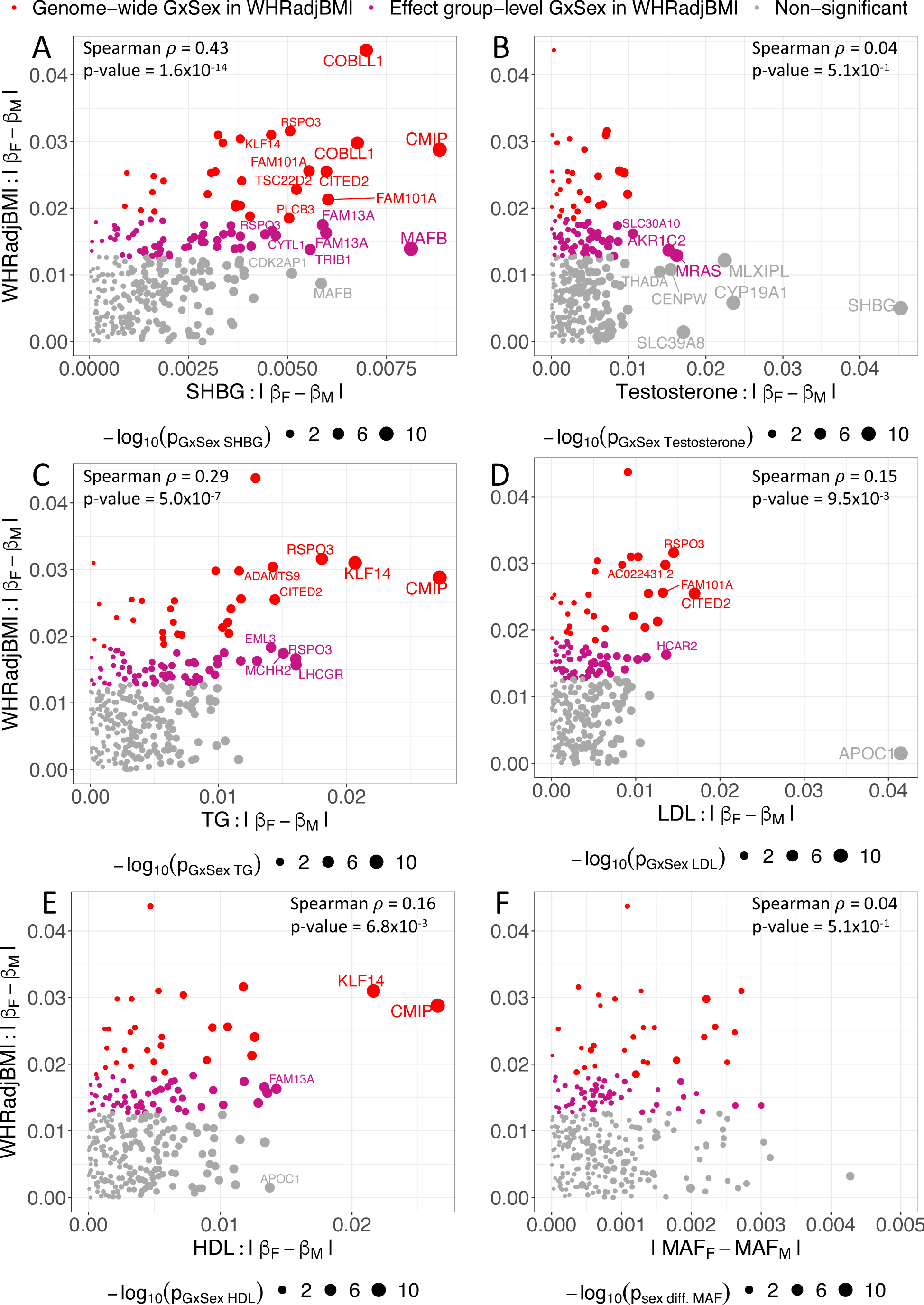
SharePro improved characterization of sex differentiated genetic effects on fat distribution. Effect size discrepancies in females and males for effect groups identified from sex stratified GWAS for waist-to-hip ratio adjusted for body mass index (WHRadjBMI) are plotted against (A) effect size discrepancies in females and males for sex hormone binding globulin (SHBG) (B) effect size discrepancies in females and males for bioavailable testosterone (C) effect size discrepancies in females and males for triglyceride (TG) (D) effect size discrepancies in females and males for low-density lipoprotein (LDL) (E) effect size discrepancies in females and males for high-density lipoprotein (HDL) (F) minor allele frequencies (MAF) discrepancies in females and males. Red dots represent genome-wide significant gene-by-sex interactions (GxSex) and violet dots represent effect group-level Bonferroni significant GxSex while grey dots represents other effect groups associated with WHRadjBMI.

In contrast, the correlation between sex differentiated genetic effects on WHRadjBMI and sex differentiated genetic effects on bioavailable testosterone was weak (Spearman correlation = 0.04; p-value = 0.51; **Figure 6B**). Since SHBG regulates both testosterone and estrogen [42], these results indicate that estrogen plays a more important role in sex differences of fat distribution. This aligns with the observation that most sex differentiated effect groups had a higher effect size in females (**Supplementary Table** S5) and supports the existing understanding that estrogen, rather than testosterone, promotes and maintains gluteofemoral fat distribution in females [36–38].

Furthermore, sex differentiated genetic effects on WHRadjBMI were more correlated with sex differentiated genetic effects on TG (Spearman correlation = 0.29; p-value = 5.0 *×* 10*^−^*^7^; **Figure 6C**) than LDL (Spearman correlation = 0.15; p-value = 9.5 *×* 10*^−^*^3^ **Figure 6D**) and HDL (Spearman correlation = 0.16; p-value = 6.8 *×* 10*^−^*^3^ **Figure 6E**). Notably, variants from effect groups demonstrating GxSex in WHRadjBMI also displayed sex differentiated effects on TG and LDL, such as those mapped to *KLF14* and *CMIP*. Interestingly, effect groups mapped to hormone receptor coding genes with important roles in the control of feeding behaviors [43–45], such as *LHCGR* and *MCHR2*, were effect group-level Bonferroni significant, which would not have been identified using the genome-wide significance threshold (**Figure 6C**).

Lastly, we did not observe sex differentiated minor allele frequencies in variants included in effect groups (**Figure 6F**), indicating limited impact of sex differentiated participation bias [26] in our study.

## 4 Discussion

In this study, we adapted SharePro, a Bayesian colocalization method [12] to propose a new framework for GxE analysis. Through extensive simulation studies, we demonstrated that accounting for effect heterogeneity can improve power for fine-mapping and by jointly fine-mapping exposure stratified GWAS summary statistics, SharePro can greatly reduce multiple testing burden in GxE analysis and achieve improved power with well-controlled FDR.

The idea of accounting for effect heterogeneity to improve power for fine-mapping is closely connected with the work by Kraft et al [18] on exploiting GxE to detect genetic associations. As illustrated in **Figure 1**, the joint approach offers advantages over the combined approach in scenarios involving effect heterogeneity, while also outperforming the stratified approach.

Furthermore, this work represents a first attempt to bridge fine-mapping and GxE analysis. In comparison to methods and study designs that screen variants based on certain assumptions to mitigate the multiple testing burden in GxE analysis [5, 7], the joint fine-mapping approach in SharePro offers the advantage of not involving additional assumptions. As a result, SharePro demonstrated robust performance across different heterogeneity settings. The SharePro approach is effective as long as the exposure stratified GWAS summary statistics are well-powered to be fine-mapped. Compared to pooling-based methods such as polygenic risk score (PRS)-based GxE analysis [46], GxE detected by SharePro has better interpretability, as PRS can involve variants associated with multiple complicated pathways.

With improved power for GxE analysis, SharePro successfully identified additional genetic effects on lung function modulated by smoking status, which were not captured by the classical heterogeneity test. In comparison to the well-characterized *CHRNA3* gene, the biological roles of the *ADAM19* and *UBR1* genes in relation to smoking are understudied, highlighting the need for further investigations, in particular, regarding their physiological roles in cellular response to smoking.

Using SharePro, we also characterized sex differentiated genetic effects on fat distribution as measured by WHRadjBMI. Our findings suggest that estrogen may play a more important role than testosterone in shaping the sex differences in fat distribution. Moreover, we observed a correlation between sex differences in fat distribution and sex differences in lipid metabolism, particularly in relation to TG levels. These results underscore the ability of SharePro in detecting biologically plausible GxE and identifying potential candidates for subsequent functional follow-up studies.

However, there are also important limitations in our study. First, the genetic effect heterogeneity detected by SharePro should be carefully interpreted based on the study design of the exposure stratified GWAS. For example, the sex differentiated genetic effects on fat distribution we identified may not reflect the actual biological differences between females and males. Instead, these differences could originated from other lifestyle factors demonstrating sex differences, such as dietary habits and physical activities [47–49]. Second, SharePro takes exposure stratified GWAS summary statistics and LD information as input and the mismatch between the LD reference panel and the true LD structure underlying samples included in GWAS can lead to convergence issue of the algorithm. We suggest using GWAS summary statistics with paired in-sample LD information to mitigate this issue. Third, in the SharePro model, we assume the LD structures are similar across different exposure categories so that the variant representations of effect groups can be shared. Further adaptations are needed for GxE analysis in crossancestry context.

In summary, we have presented SharePro to account for effect heterogeneity in fine-mapping and improve the power of GxE analysis. We envision this new analytical framework will have important utility in characterizing genetic effects heterogeneity in complex traits and diseases and identifying modifiable risk factors for prevention and management of complex diseases.

## 5 Data and Software Availability

The SharePro software for detecting GxE is openly available at https://github.com/zhwm/SharePro_gxe and the analysis conducted in this study is available at https://github.com/zhwm/SharePro_gxe_analysis. Individual-level phenotype and genotype data from the UK Biobank are available upon successful application at https://www.ukbiobank.ac.uk. GEM was obtained from GitHub at https://github.com/large-scale-gxe-methods/GEM and PLINK was obtained from https://www.cog-genomics.org/plink/.

## 6 Supporting Information

### 6.1 Supplementary Tables

### 6.2 Supplementary Figures

## Acknowledgements

W.Z. has been supported by a doctoral training fellowship from the FRQNT (319188) and the Healthy Brains, Healthy Lives Program, funded by the Canada First Research Excellence Fund (CFREF), Quebec’s Ministère de l’Économie et de l’Innovation (MEI), and the Fonds de recherche du Québec (FRQS, FRQSC and FRQNT). Y.L. is supported by Natural Sciences and Engineering Research Council (NSERC) Discovery Grant (RGPIN-2019-0621), Fonds de recherche Nature et technologies (FRQNT) New Career (NC-268592), and Canada First Research Excellence Fund Healthy Brains for Healthy Life (HBHL) initiative New Investigator start-up award (G249591). H.S.N holds a Canada Research Chair funded by the Canadian Institutes of Health Research and has been supported by NSERC Discovery Grant (RGPIN-2018-05962). This research used the NeuroHub infrastructure and was undertaken thanks in part to funding from the Canada First Research Excellence Fund, awarded through the Healthy Brains, Healthy Lives initiative at McGill University. This research was enabled in part by support provided by Calcul Québec and the Digital Research Alliance of Canada. This research has been conducted using the UK Biobank Resource under Application Number 45551.

## 9 Author contributions

Conceptualization: W.Z.; Data curation: W.Z.; Formal analysis: W.Z.; Funding Acquisition: W.Z., Y.L., H.S.N and J.D.; Investigation: W.Z.; Methodology: W.Z.; Project Administration: R.S., H.S.N and J.D.; Resources: Y.L., H.S.N and J.D.; Software: W.Z.; Supervision: Y.L., R.S., H.S.N and J.D.; Validation: W.Z., R.S., Y.L., H.S.N and J.D.; Visualization: W.Z.; Writing – Original Draft Preparation: W.Z.; Writing – Review & Editing: W.Z., R.S., Y.L., H.S.N and J.D.

## 10 Disclosures

The authors declare no conflict of interest.

## Notes

https://github.com/zhwm/SharePro_gxe

https://github.com/zhwm/SharePro_gxe_analysis

### Competing Interest Statement

The authors have declared no competing interest.

