## Supplementary Figure for "Accounting for genetic effect heterogeneity in fine-mapping and improving power to detect gene-environment interactions with SharePro"

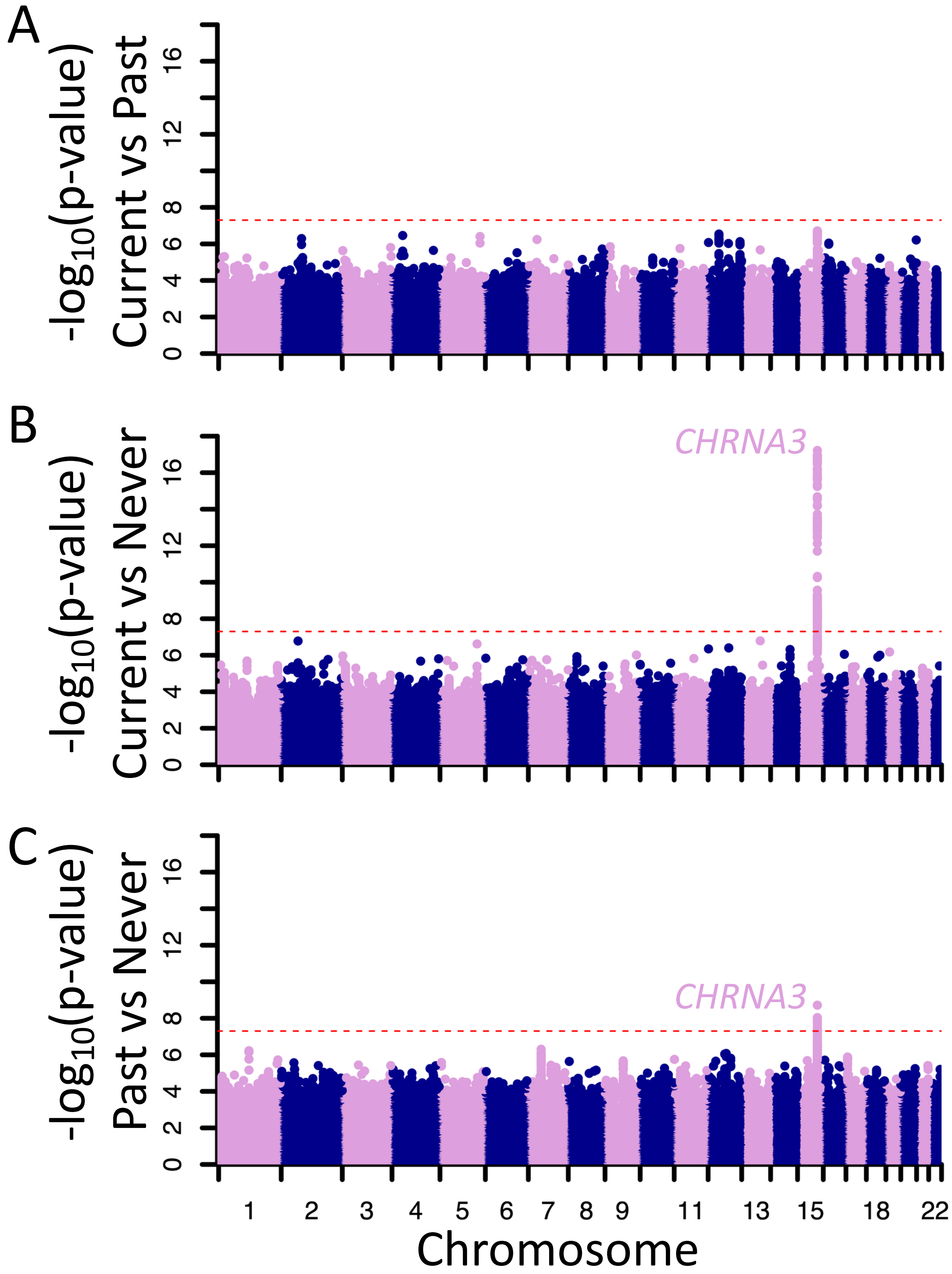


**Supplementary Figure S1: GxSmoking detected using the heterogeneity test.** (A) Variant-level GxSmoking p-values comparing current smokers with past smokers are plotted against the genomic positions of variants. (B) Variant-level GxSmoking p-values comparing current smokers with never smokers are plotted against the genomic positions of variants. (C) Variant-level GxSmoking p-values comparing past smokers with never smokers are plotted against the genomic positions of variants. The red dotted line corresponds to a genome-wide significance threshold of 5e-8.
